# Evaluating and mitigating clinical samples matrix effects on TX-TL cell-free performance

**DOI:** 10.1101/2022.05.02.489947

**Authors:** Peter L. Voyvodic, Ismael Conejero, Khouloud Mesmoudi, Eric Renard, Philippe Courtet, Diego I. Cattoni, Jerome Bonnet

**Affiliations:** Centre de Biologie Structurale (CBS). INSERM U1054, CNRS UMR5048, University of Montpellier, France; Department of Psychiatry, CHU Nîmes, University of Montpellier, Nîmes, France; UMR CNRS 5203/INSERM U1191, Institute of Functional Genomics, University of Montpellier, Montpellier, 34090, France; Department of Endocrinology, Diabetes, Nutrition and CIC INSERM 1411, Montpellier University Hospital, Montpellier, 34295, France; CHU Montpellier, Department of Emergency Psychiatry and Post Acute Care, Lapeyronie Hospital, CHRU Montpellier, Montpellier, 34090, France

**Keywords:** synthetic biology, cell-free, biosensor, diagnostics, clinical samples

## Abstract

Cell-free biosensors are promising tools for medical diagnostics, yet their performance can be affected by matrix effects arising from the sample itself or from external components. Here we systematically evaluate the performance and robustness of cell-free systems in serum, plasma, urine, and saliva using two reporter systems, sGFP and luciferase. In all cases, clinical samples have a strong inhibitory effect. Of different inhibitors, only the RNase inhibitor mitigated matrix effects. However, we found that the recovery potential of RNase inhibitor was partially muted by interference from glycerol contained in the commercial buffer. We solved this issue by designing a strain producing an RNase inhibitor protein requiring no additional step in extract preparation. Furthermore, our new extract yielded higher reporter levels than previous conditions and tempered interpatient variability associated with matrix effects. This systematic evaluation and improvements of cell-free system robustness unified across many types of clinical samples is a significant step towards developing cell-free diagnostics for a wide range of conditions.

## INTRODUCTION

Biosensors are detection tools that integrate a biological recognition element from a sensor module and transduce the signal into a quick, measurable response^1^. Rapid point-of-care testing applications are an alternative to complex technical procedures in clinical settings and reduce the need for time-consuming and equipment-dependent sample processing. Hence, portable biosensors allow close monitoring of chronic disease, as exemplified by portable glucose monitoring devices that have revolutionized diabetes care^2^. These tools are adapted to a real-time assessment of such clinical populations and more precisely determine clinical outcomes. In this way, less biased data are simply and repeatedly sampled outside the clinical setting while taking into account the patient’s own environment and variability^3^. Moreover, biosensors are particularly promising tools for field diagnostics in low-resource settings^4^, especially in the context of transmissible infectious and endemic diseases such as HIV^5^, malaria^6^, or Zika^7^. The poor testing response of many countries to COVID-19 has highlighted the importance of rapid, low-cost, and easily distributable diagnostic devices worldwide^8^.

Among the vast array of available technologies, cell-free expression systems have recently emerged as promising candidates for versatile biosensor engineering^9,10^ as they support the operation of sophisticated genetic circuits while requiring small reaction volumes^11^. Compared to whole-cell biosensors, cell-free systems can detect molecules like nucleic acids that do not cross cellular membranes, as well as ones typically toxic to living cells. In addition to being abiotic and non-replicating, with little need for biocontainment measures, cell-free expression systems endure no evolutionary pressure that can alter whole-cell biosensors. Reactions can occur at ambient or body temperature, by taping the sensor to the skin for example^12^, eliminating equipment like incubators at the point of detection. Performances are easily tunable by varying extract and plasmid composition, and the protein expression yield may be optimized through a large variety of methods^13^. Cell-free biosensors can be lyophilized and kept stable at room temperature for up to one year^14^ and may provide rapid responses in as little as under one hour^15^. The combination of engineered cell-free transcription and translation (TX-TL) systems with electrochemical platforms further enables multiplexed biomarker detection^16^.

Overall, with short reaction times, long-term storage, and the compatibility with nano-electrode interfaces^16^, cell-free systems are good candidates toward point-of-care testing of pathological biomarkers in clinical settings and remote locations. The emergence of decentralized, portable diagnostics may improve the acquisition of epidemiological data worldwide, which is proving critical in planning global healthcare measures to mitigate the effects of non-communicable diseases and recent outbreaks, such as Zika or the COVID-19 pandemic^17^. Yet, some issues are currently limiting the translation of cell-free diagnostic platforms to clinical use, such as cross detection of non-identified markers, or most importantly, the interference from biological samples. Variability in the components of complex samples can affect the readouts of even sophisticated analytical instrumentation, a phenomenon known as matrix effects that can yield inaccurate results^18^. These effects were reported in several types of biosensors produced in *E. coli* extracts that have been tested using human samples (**Table 1**).

**Table 1:** Summary of previous cell-free biosensors testing in clinical samples, using *E. coli* extracts.

| Detected marker | Clinical sample | Optimization | Clinical condition | Ref. |
| --- | --- | --- | --- | --- |
| Spiked cocaine and endogenous hippuric acid | Urine | RNase Inhibitor | Hospitalized patients | 15 |
| Spiked BPA, E2, and DPN | Whole Blood and Urine | RNase Inhibitor | Healthy donors | 19 |
| 3-oxo-C12-HSL | Sputum | Three extract strains | *P. aeruginosa*-infected patients | 20 |
| Spiked TRIAC and T3 hormones | Whole Blood and Urine | -None- | Human volunteers | 21 |
| Zinc | Serum | RNase Inhibitor | Human volunteers | 18 |
| Glutamine | Saliva | RNase Inhibitor | Human volunteers | 22,23 |

Although the final aim of cell-free biosensors is their field deployment and the point-of-care testing of pathogenic biomarkers, only sparse studies have systematically assessed the performance of cell-free biosensors in multiple different types of human biological samples^19,22,23^. Here, we monitored the performance of *E. coli* TX-TL cell-free extract across four types of clinical samples taken from patients in hospital settings through minimally invasive methods. Samples were not processed before testing except in their basic preparation (*i*.*e*. blood centrifugation for serum and plasma after collection in appropriate vacuum tubes). We tested matrix effects of the samples on both constitutively-produced superfolder Green Fluorescent Protein (sfGFP) and firefly luciferase^15^. In addition, we systematically examined the improvement of the *E. coli* TX-TL cell-free system by adding RNase and protease inhibitors. Importantly, we found that the glycerol present in commercial RNase inhibitors reduces cell-free production. As such, we developed an *E. coli* strain that can produce its own RNase inhibitor *in situ* during extract production without any additional procedure steps, simultaneously eliminating the cost of commercial inhibitors and increasing protein production in the presence of clinical samples over that obtained when they are used. Finally, we examined the interpatient variability effects through testing ten serum, plasma, and urine patient samples. Our new extract reduced interpatient variability, particularly for plasma samples. By probing the effects of both pooled and individual patient samples across an array of different types of clinical samples and providing an easier way to mitigate their matrix effects, our results represent a significant step forward in developing cell-free diagnostics for a range of biomarkers and pathological conditions.

## RESULTS

### Clinical sample matrix strongly inhibits reporter production in cell-free biosensors

We first measured the matrix effects of various clinical samples (serum, plasma, urine, and saliva) on cell-free activity. To do so, we monitored the production of two constitutively expressed reporters, superfolder GFP (sfGFP) and firefly luciferase (Luc) in the presence or absence of clinical samples. We chose sfGFP and luciferase because they are both common reporters widely used for signal quantification. Plasmids constitutively expressing sfGFP or luciferase were mixed with *E. coli* TX-TL extract prepared using a French press as previously described^15^, and an optimized buffer containing the necessary building blocks, salts, and energy source for transcription and translation. Finally, as these core reaction components take up 80-90% of the available reaction volume, non-processed clinical samples were added to the reaction mix as 10% of the final reaction volume (**Figure 1A**).

**Figure 1:**
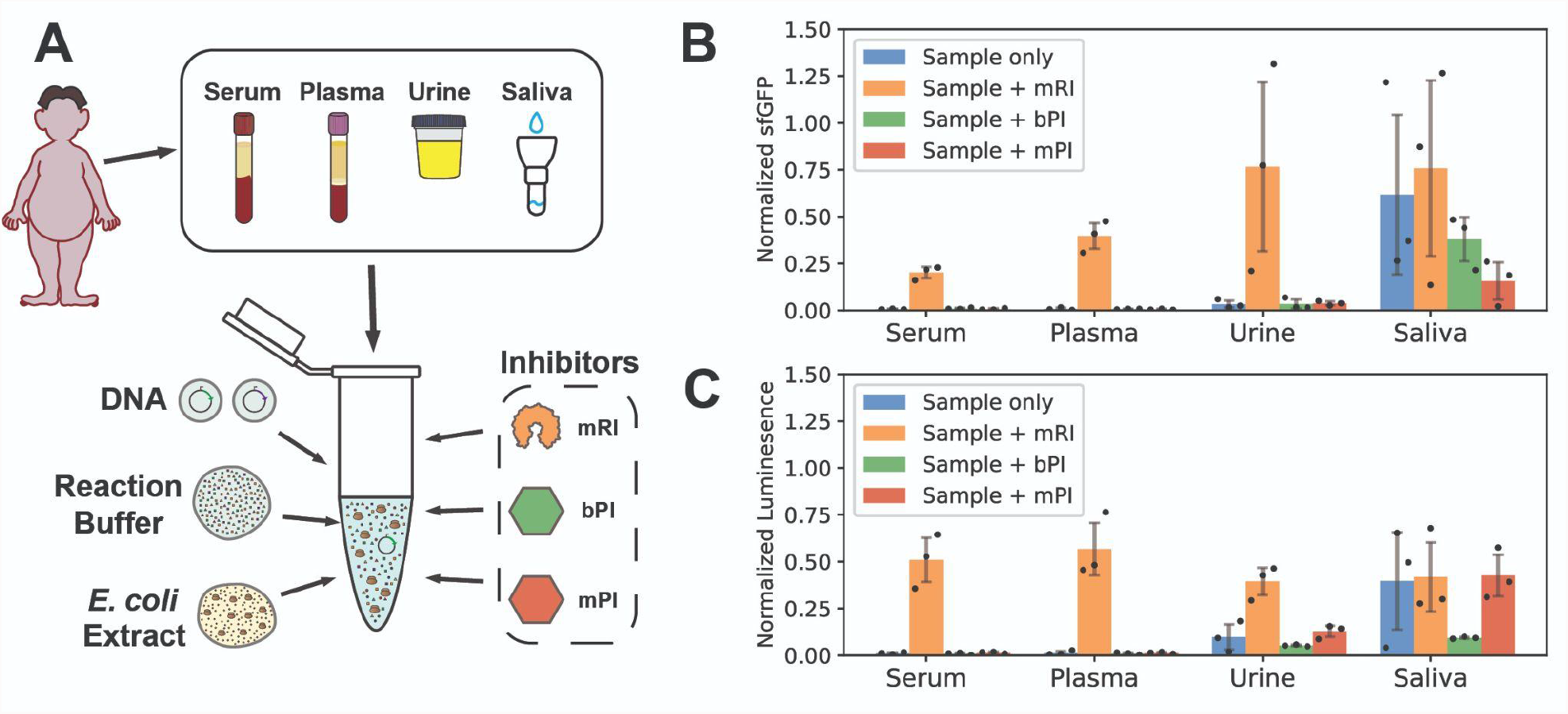
Deleterious clinical samples matrix effect on reporter expression is partially recovered by commercial RNase inhibitor. **A**. Plasmids encoding sfGFP and luciferase proteins were mixed with *E. coli* cell extracts, optimized reaction buffer, and clinical samples (serum, plasma, urine, or saliva). **B-C**. Quantification of the matrix effects. 100 nM of constitutively expressing sfGFP (B) or luciferase (C) plasmids was added to a cell-free system containing 10% clinical sample with or without murine RNase inhibitor (mRI), bacterial protease inhibitor (pI), or mammalian protease inhibitor (mPI). All measurements were normalized relative to the positive control (without clinical sample or inhibitors). Each clinical sample is a pool from three different individuals. Data are the mean of three experiments performed on three different days. Error bars correspond to ±SD.

We quantified the matrix effects of different clinical samples on constitutive reporter expression in the absence or presence of RNase inhibitor and two protease inhibitors (bacterial and mammalian) relative to a positive control with neither clinical sample nor inhibitor (**Figure 1B-C**). We found that all clinical samples had an inhibitory effect on reporter production, albeit to a different extent. Without any inhibitors, serum and plasma almost completely impeded reporter production (>98% inhibition with respect to no sample addition). Urine inhibited more than 90% of reporter production for both sfGFP and Luc with respect to no sample addition, whereas saliva produced the least interference for both reporters (70% inhibition on luciferase and 40% on sfGFP, with respect to control).

### Commercial RNase inhibitor partially restores cell-free activity while protease inhibitors display poor mitigation of matrix effects

We tested two categories of additives inhibiting enzymatic activities that could affect the reactions: RNases and proteases. RNAse inhibitor was previously shown to improve the efficiency of cell-free reactions^19^ in some types of clinical samples or were systematically added to cell-free reactions with biological fluids^15,18^. Also, endogenous proteases from *E. coli* extracts have been suggested to affect the yield of protein synthesis^24,25^. We choose to test both bacterial and mammalian protease inhibitors to account for proteases found in both the *E. coli* extract and in the human clinical samples. While the use of RNase inhibitors has been previously tested^19^, no previous study has evaluated neither protease inhibitors nor all four of the clinical samples described here.

The addition of RNase inhibitor improved sfGFP production by about 70% in urine, 20% in serum, and 40 % in plasma, while bacterial and mammalian protease inhibition failed to improve cell-free reaction performance in any of the clinical samples (**Figure 1B**). Results were comparable in all conditions when using the firefly luciferase reporter. The addition of RNase inhibitor restored luciferase signals in saliva, plasma, serum, and to a lesser extent in urine, reaching 50% of the luciferase production from the absence of a clinical sample (**Figure 1C**). As with sfGFP, bacterial and mammalian protease inhibitors did not produce significant improvement on cell-free protein synthesis for the Luc reporter.

### Glycerol present in the commercial buffer of enzymatic inhibitors is responsible for signal degradation in cell-free reactions

While RNase inhibitors generally led to increased protein production in the presence of clinical samples, in all cases full signal (i.e. when no clinical sample was present) was never recovered. Additionally, we tested the effect of inhibitors in cell-free reactions without any clinical sample (**Supplementary Figure 1**) and observed that all of them degraded reporter production, with RNase inhibitor being the most detrimental (∼50% reduction of signal with respect to no inhibitor). We wondered if the buffer composition of the commercial RNAse inhibitor could be responsible for this phenomenon. First, we compared cell-free reaction efficiency, with and without RNase inhibitor, vs. buffer with identical components as listed by the manufacturer (50 mM KCl, 20 mM HEPES, 8 mM DTT, 50% glycerol) (**Figure 2A**). Indeed, there was a marked decrease in protein production from adding RNase inhibitor that was identical to that of adding the buffer alone. This result was consistent over a range of plasmid concentrations (**Supplementary Figure 2**). To disentangle which part of the buffer was responsible, we separately added each component of the commercial buffer to the cell-free reaction, individually and in all possible combinations (**Figure 2B**). We found that glycerol alone (at 1% final reaction concentration) accounted in all cases for the sfGFP production decrease independently of the presence of any other component of the buffer. These data demonstrate that the decrease of cell-free reaction performance observed when adding RNase inhibitor is exclusively due to the glycerol contained in the buffer solution.

**Figure 2:**
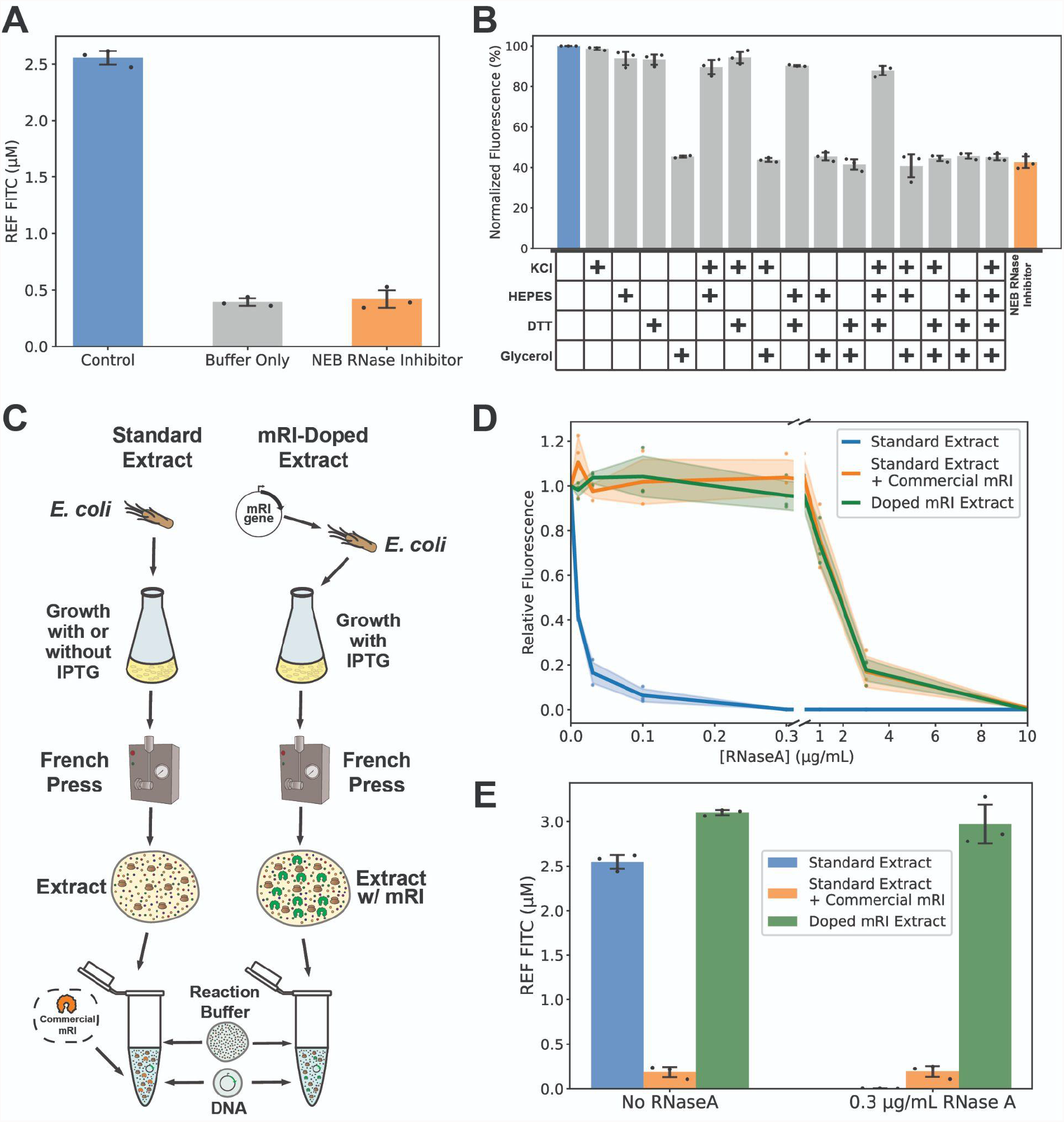
*In situ* RNAse production recovers reporter production originally thwarted by glycerol. **A**. sfGFP expression for the positive control (standard cell-free reaction mix) and in the presence of RNase inhibitor (mRI) or equivalent buffer concentration without mRI (50 mM KCl, 20 mM HEPES, 8 mM DTT, 50% glycerol). **B**. Effect of individual and combined commercial buffer components on reporter output. **C**. Scheme of cell-free extract production and reaction mix. mRI-doped extract was obtained by cloning the murine RNase inhibitor gene under T7 promoter control. Purification and reaction preparation were equivalent to standard extract, but no additional mRI was required to perform cell-free measurements. **D**. Relative expression of sfGFP in standard extract (BL21 Star), standard extract with commercial RNase inhibitor (BL21 Star + commercial mRI), and extract induced to produce RNase inhibitor during cell growth (BL21 Star + Doped mRI Extract) when challenged with an RNaseA concentration gradient. **E**. Fluorescent intensity (relative equivalence of µM FITC) of standard extract with and without commercial mRI vs. mRI-doped extract at 0 and 0.3 µg/mL RNaseA. Data in Figure 2A are the mean of three technical replicates. Data in Figures 2B, 2D, and 2E are the mean of three experiments performed on three different days. Error bars or shaded areas correspond to ±SD.

### In situ RNase inhibitor production allows equivalent protection to that of commercial inhibitor while allowing higher reporter production

We then hypothesized that we could avoid glycerol inhibition by expressing RNase inhibitor in *E. coli* while growing the cell-free extract. This would prevent the need for glycerol and have the added benefit of reducing the overall reaction cost. After cloning a codon-optimized version of the murine RNase inhibitor (mRI) gene into a plasmid under a T7 promoter, we transformed it into competent *E. coli* for extract production. Because adding IPTG during the growth process is a common cell-free practice for producing T7 RNA polymerase *in situ*, having the mRI gene under a T7 promoter would allow both high mRI production and potentially require no additional steps in the process (**Figure 2C**).

To examine whether our mRI-doped strain could indeed protect against RNases as efficiently as the commercial product, we evaluated reporter signal vs. increasing concentrations of RNaseA for the standard extract with and without commercial mRI against our mRI-doped extract (**Figure 2D**). While protein production with no inhibitor sharply drops to almost no production with less than 0.1 µg/mL RNaseA, both commercial and doped mRI extracts show no decrease in signal up to 0.3 µg/mL of RNaseA and can still produce a measurable signal when confronted with up to 3 µg/mL. Importantly, when examining the absolute signal of each condition, the mRI-doped extract shows more than ten times higher levels of protein production than the glycerol-inhibited commercial mRI reactions (**Figure 2E**). Thus, our mRI-doped extract provided equivalent levels of relative RNase protection to that of the commercial inhibitor while allowing for much higher absolute levels of protein production.

### RNase inhibitor-doped cell extracts outperform the use of commercial reagents with clinical samples

We sought to compare the performance in clinical samples of our doped extract relative to extract supplemented with commercial RNAse inhibitor. We tested serum, plasma, urine, and saliva using standard extract, standard extract with commercial mRI, and doped extract for constitutive sfGFP expression (**Figure 3A-B**). While matrix effects still reduced overall protein expression, the doped extract showed marked improvement in fluorescent signal over standard extract supplemented with commercial inhibitor (**Figure 3C**). This demonstrates that the effects observed in reducing RNase activity observed with extract-expressed RNase translate to an improvement with clinical samples as well, while avoiding the costs and glycerol effects of commercial inhibitors.

**Figure 3:**
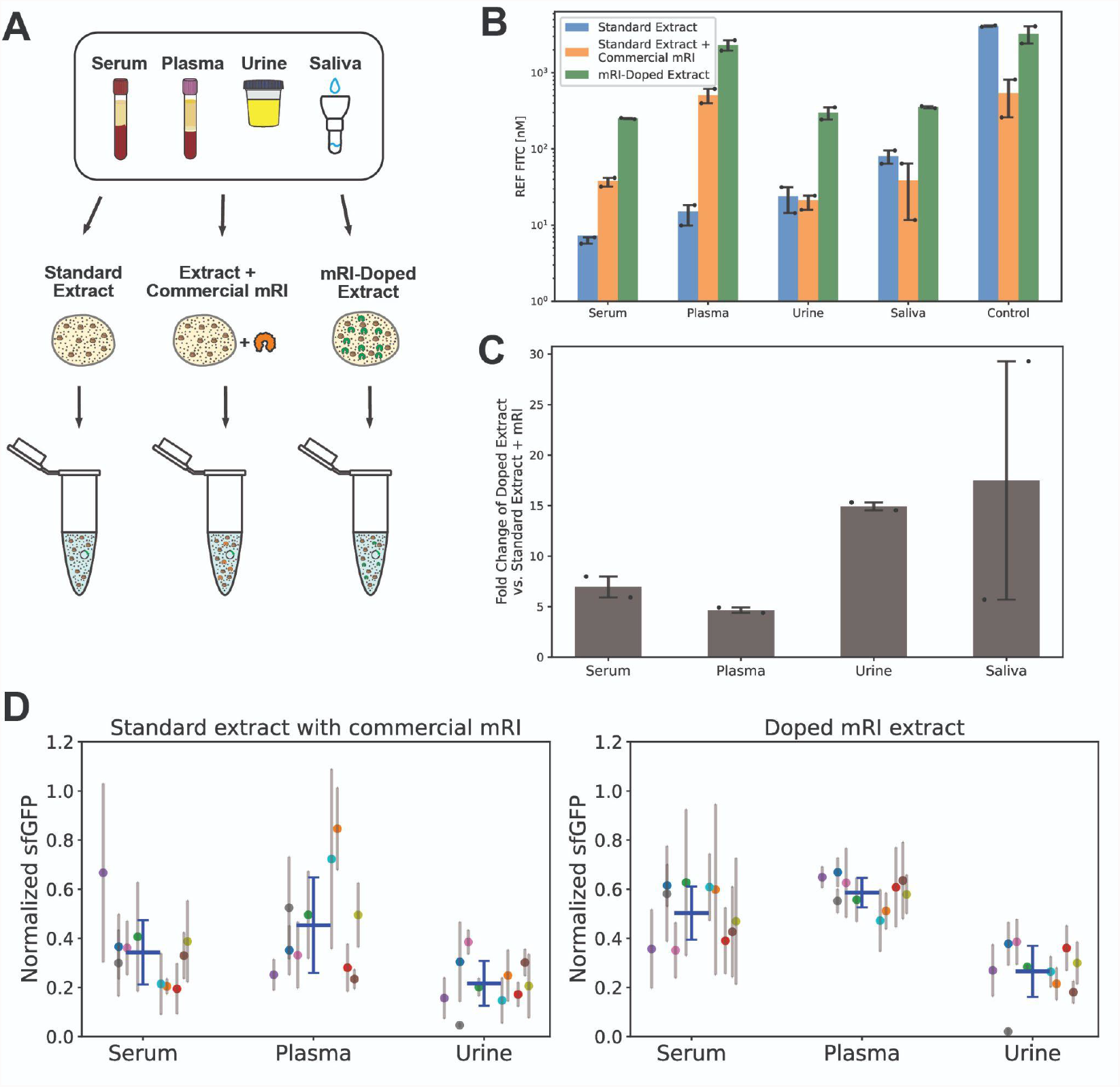
Doped mRI extract yields larger protein production and moderates interpatient matrix effects. **A**. Clinical samples (serum, plasma, urine) were mixed with buffer, 10 nM sfGFP DNA, and one of three *E. coli* extract conditions: standard extract, standard extract with commercial mRI, or doped extract with *in situ*-produced mRI. **B**. Quantification of the matrix effects for each sample. Fluorescence intensity was normalized to the equivalent concentration of FITC (nM). **C**. Fold change in sGFP production of mRI-doped vs. standard extract with commercial mRI. Each clinical sample is a pool of three different individuals. Data are the mean of two experiments performed on two different days. Error bars correspond to ±SD. **D**. Individual patients’ sGFP production for each clinical sample. Samples from ten patients were measured independently on three different days. Colored solid dots and thin gray lines indicate the mean and SD for each patient. Solid blue lines indicate the mean and SD for the population.

Most studies examining the development and optimization of cell-free based biosensors employ pooled clinical samples or artificial equivalents of clinical samples. We thought that our system, which did not directly measure the presence of any biomarker, represented an optimal benchmark opportunity to study matrix variability from patient to patient. Furthermore, we wished to test if *in situ* production of mRI further mitigated interpatient matrix effects. We examined, over three consecutive days, the ten different patient samples of serum, plasma, and urine, ensuring independence between measurements (**Figure 3D**) on the presence of commercial inhibitors or with our mRi doped extract. In all cases, the dispersion of the data was lower when employing doped mRI extract, particularly for plasma and urine. These results highlight that despite improved protein production with RNase inhibitors, significant challenges likely remain to develop cell-free diagnostics with sufficient robustness to function with the variability between patient samples and sampling conditions in a point-of-care setting.

## DISCUSSION

Our workflow provides unified data and systematic studies of the performance of a TX-TL cell-free biosensor in a wide variety of clinical human samples. We studied and compared the matrix effects in complex media and assessed simple and reproducible optimization strategies. We further examined an expanded range of inhibitors likely to have a beneficial effect on sensor performance, including mammalian and protease inhibitors. The knowledge of the interference with human biological samples and of possible optimization strategies is important to translate the use of cell-free based biosensors to real clinical populations with a broad range of pathological conditions. This step is crucial to achieve their potential as next-generation platforms for rapid, low-cost, field diagnostics. Our procedure is straightforward, reproducible, and may be applied in real clinical settings to validate the use of TX-TL cell-free biosensors on a large scale.

One of the strengths of our workflow is the absence of sample preprocessing that would require heavy technical methods, such as analyte extraction or protein phase separation^20^. The only sample processing was the standard centrifugation procedure applied to obtain serum and plasma from whole blood tubes. Our data support the expansion of cell-free biosensors to a variety of fresh human samples in remote settings without the need for complex technical sample processing. In assays where blood plasma or serum are required, these biosensors could be coupled with low-tech centrifuges, like the paperfuge or egg-beater centrifuge, for sample preparation^26,27^.

We compared the signal output from two well-known reporter modules, sfGFP and luciferase, in a variety of clinical samples. The reporter’s signal was strongly thwarted by the addition of clinical samples in all cases, although saliva showed the least inhibition effects. The significant drop in performance and partial restoration by addition of RNAase inhibitor in serum and plasma may be explained by the large amounts of pancreatic type ribonucleases secreted by endothelial cells from arteries, veins, and capillaries^28^. Similarly, human urine has long been known to contain multiple types of RNase activity^29^. Interestingly, while human saliva can exhibit ribonuclease activity, particularly under certain pathological conditions^30^, our results suggest that it does not itself always significantly contribute to the sample matrix effects. One potential downside to sfGFP as a reporter for point-of-care diagnostics is the need to produce low-cost devices that can excite the fluorescent protein and filter out the emission signal; however, other groups have worked to design 3D printed fluorescent readers that could work well for this purpose^31^. Compared with sfGFP use, the drawback to using luciferase in clinical settings is the need to add luciferase substrate (luciferin) to trigger light emission. However, luciferase-associated sensors have the advantage of providing a detectable signal for simple optical devices that need no external light source for stimulation. Some studies have reported simple signal detection with smartphone cameras which may be easily handled at the patient’s bedside^32,33^. Additionally, other reporters that induce a color change, such as LacZ^14^ or C23DO^12^, could provide a simple, visual output result.

Some of the specific effects of the RNase inhibitor were likely muted, impairing full signal recovery. We found that glycerol contained in the RNase inhibitor commercial buffer was responsible for the reduced efficiency of cell-free reactions. While evidence of glycerol inhibition of cell-free reactions has been previously reported^34^, the lack of current glycerol-free RNase inhibitors precludes another viable commercial option. We decided to sidestep the glycerol effects by designing an *E. coli* plasmid that could produce mRI *in situ* during the extract preparation process. We found that by producing mRI under the control of a T7 promoter during the cell growth phase, we created a cell extract that could protect against RNaseA as well as a commercial RNase inhibitor without the overall signal reducing effects of buffer glycerol. Additionally, due to the absence of glycerol, this extract exhibited much higher signal in the presence of clinical samples. While recent publications have similarly produced mRI to help protect against clinical matrix effects, they required multiple reactions or the combination of multiple extracts grown at different temperatures with additional folding chaperones ^22,23^. In contrast, our strain does not require any additional steps, aside from adding IPTG during cell growth, which is already commonly done to induce T7 polymerase production in cell-free extracts.

Finally, we examined the interpatient variability of three different clinical samples: serum, plasma, and urine using a commercial inhibitor and our doped mRI extract. Using ten individual samples of each, we found that the matrix effects produced less variability between patients when using the doped extract, particularly for plasma and urine. This kind of validation, using a large number of individual patients, is essential for the development of new biosensors, as we show that matrix effects can be so significant as to completely abolish protein production in one patient’s urine sample. As concentrations of different ions and metabolites can vary widely in urine depending on patient hydration^35^, it is perhaps not surprising to see large variability; however, these findings highlight that significant robustness remains to be engineered into cell-free diagnostics before they can be reliably used in a point-of-care setting. Interestingly, blood plasma showed the lowest interpatient variability. While blood plasma and serum can frequently be used interchangeably, the coagulation process in acquiring serum can lead to changes in certain platelet components, such as aspartate aminotransferase, lactate dehydrogenase, fibrin/fibrinogen, and serotonin^36^. Further studies could investigate which components specifically influence cell-free expression, but our work indicates that plasma may be a good clinical sample candidate for future applications requiring a high degree of reproducibility as in chronic conditions or long-term follow-up studies or diagnostics.

Overall, these results represent a comprehensive overview of some of the promise and potential challenges of using cell-free systems for clinical diagnostics with unprocessed samples. In addition to providing a dataset that compares four types of commonly used clinical samples across a unified set of experiments, it represents, to the best of our knowledge, the first time that blood plasma has been used in cell-free reactions. Additionally, saliva is of particular interest since it involves a pain-free, non-invasive collection process and can be potentially used to monitor the advancement of neurodegenerative and neuropsychiatric disorders like autism, Alzheimer’s, Parkinson’s, and Huntington’s disease, in addition to the oral microbiome, which is involved in dental and periodontal health^37,38^. Opening cell-free biosensors to a broader range of clinical samples and ensuring their reliability through systematic optimization will help provide portable, low-cost diagnostics tools to address global health challenges.

## MATERIALS AND METHODS

### Molecular biology

The reporter plasmid for sfGFP (pBEAST2-sfGFP) was previously characterized^15^ and is available on Addgene (#126575). The plasmid for luciferase (pBEST-Luc) was obtained from Promega. DNA for cell-free reactions was prepared from overnight bacterial cultures using Maxiprep kits (Macherey-Nagel).

To create the plasmid to induce murine RNase inhibitor expression in *E. coli* extract, the amino acid sequence was obtained from Uniprot (#Q91VI7) and codon-optimized for *E. coli* and synthesized by Twist Bioscience. It was then cloned by Gibson assembly into the pBEAST backbone under the control of a T7 promoter and transformed into either BL21 Star or BL21 Star (DE3)::RF1-CBD3 *E. coli* for extract production. The final plasmid, pPLV_C1, will be available from Addgene.

### Cell-extract preparation

Cell-free *E. coli* extracts were produced using a modified version of existing protocols as previously detailed^15^. Briefly, an overnight culture of BL21 Star or BL21 Star (DE3)::RF1-CBD3 *E. coli* was used to inoculate 660mL of 2xYT-P media in each of six 2 L flasks at a dilution of 1:100. The cultures were grown at 37 °C with 220 r.p.m. shaking for ∼3.5 h until OD 600 = 2.0. For the mRI-doped extract, cultures were induced with 1 mM IPTG at OD 600 = 0.4-0.6. Cultures were spun down at 5000 × g at 4 °C for 12 min. Cell pellets were washed twice with 200 mL S30A buffer (14mM Mg-glutamate, 60mM K-glutamate, 50mM Tris, pH 7.7), centrifuging afterward at 5000 × g at 4 °C for 12 min. Cell pellets were then resuspended in 40mL S30A buffer and transferred to preweighed 50 mL Falcon conical tubes, where they were centrifuged twice at 2000 × g at 4 °C for 8 and 2 min, removing the supernatant after each. Finally, the tubes were reweighed and flash-frozen in liquid nitrogen before storing at −80 °C.

Cell pellets were thawed on ice and resuspended in 1 mL S30A buffer per gram cell pellet. The suspensions were lysed via a single pass through a French press homogenizer (Avestin; Emulsiflex-C3) at 15,000–20,000 psi and then centrifuged at 12,000 × g at 4 °C for 30 min to separate out cellular cytoplasm. After centrifugation, the supernatant was collected and incubated at 37 °C with 220 rpm shaking for 60 min to digest the remaining mRNA with endogenous nucleases. Subsequently, the extract was re-centrifuged at 12,000 × g at 4 °C for 30 min and the supernatant was transferred to 12–14 kDa molecular weight cut-off (MWCO) dialysis tubing (Spectrum Labs; Spectra/Por4) and dialyzed against 2 L of S30B buffer (14 mM Mg-glutamate, 60 mM K-glutamate, ∼5mM Tris, pH 8.2) overnight at 4 °C. The following day, the extract was re-centrifuged at 12,000 × g at 4 °C for 30 min. The supernatant was aliquoted and flash-frozen in liquid nitrogen before storage at −80 °C.

### Cell-free sensor reaction optimization

Cell-free reactions were prepared by mixing 33.3% cell extract, 41.7% buffer, and 25% plasmid DNA, clinical sample, inhibitors, and water. Buffer composition was such that final reaction concentrations were as follows: 1.5 mM each amino acid, 50 mM HEPES, 1.5 mM ATP and GTP, 0.9 mM CTP and UTP, 0.2 mg/mL tRNA, 0.26 mM CoA, 0.33 mM NAD, 0.75 mM cAMP, 0.068 mM folinic acid, 1 mM spermidine, 30 mM 3-PGA, and 2% PEG-8000. In addition, the Mg-glutamate (0–20 mM), K-glutamate (20–300 mM), and DTT (0–3 mM) levels were serially calibrated for each batch of cell extract for maximum signal. Reactions using constitutively expressing sfGFP or luciferase plasmids were performed at a final DNA concentration of 10 nM or 100 nM. For testing with an RNaseA gradient, RNaseA (Qiagen) was diluted in water and added at 10% reaction volume to their final concentrations.

The sfGFP-output reactions were prepared in PCR tubes on ice and 20 μL was transferred to a black, clear-bottom 384-well plate (ThermoScientific), sealed, and the reaction was carried out at 37°C in a plate reader (Biotek; Cytation3) to measure fluorescence over time. Mean equivalent fluorescence (MEF) of soluble fluorescein isothiocyanate (FITC) was calculated based on measurements of a standard gradient under identical excitation/emission and gain settings as previously published by ourselves and others.^31,39^ Luciferase-output reactions were incubated at 37°C for 8 hours. Samples were then transferred to white 96-well plates and 50 μL of Bright-Glo Luciferase Assay Reagent (Promega) was added and mixed by manual orbital agitation. The plates were sealed and luciferase levels were measured in a plate reader 5 min after the addition of the reagent. The subsequent data were processed and graphs were created using Python scripts.

### Cell-free reactions with human urine, saliva, plasma, and serum

Cell extract and buffer conditions were maintained from those used in optimization reactions. Saliva, urine, serum, and plasma were pooled from three individual patient samples to reduce patient-specific inhibition effects. 10% reaction volume of each sample was included in 20 μL reactions containing extract, buffer, DNA, and inhibitors when applicable. Human urine samples were obtained from the Endocrinology Department at the University of Montpellier in accordance with ethics committee approval (#190102). Human blood samples were obtained from the EFS (Etablissement Français du Sang). Saliva was issued from the samples collected in the SADS-CS study conducted in the Department of Emergency Psychiatry and Acute Care, Hôpital Lapeyronie, CHU Montpellier (ethics committee approval: CPP Sud-Méditerranée-IV). In addition, where applicable reactions were supplemented with 0.32 units of murine RNase Inhibitor (New England Biolabs, #M0314), 1X equivalent of mammalian protease inhibitor (100X stock solution Euromedex, #B14011), or 1X equivalent of bacterial protease inhibitor (10X stock solution Sigma-Aldrich, #P8465).

## ACKNOWLEDGEMENTS

We thank the Bonnet lab members for fruitful discussions. We are grateful to Pau Bernado, Annika Urbanek, and Anna Morato for helpful discussions on cell-free systems and extract creation, Gottfried Otting for BL21 (DE3) Star::RF1-CBD3 strain, and the patients for providing their samples. This work was supported by an ERC starting grant “COMPUCELL” and ANR “SynbioDiag” to J.B. J.B. also acknowledges continuous support from the INSERM Atip-Avenir program and the Bettencourt-Schueller Foundation. The CBS acknowledges support from the French Infrastructure for Integrated Structural Biology (FRISBI) ANR-10-INSB-05-01. All plasmids will be available from Addgene.

## AUTHOR CONTRIBUTIONS

P.L.V., I.C., D.I.C., and J.B. designed experiments. P.L.V, I.C. and K.M. performed experiments. I.C., E.R., and P.C. participated in clinical sample collection. P.L.V., I.C., D.I.C., and J.B. analyzed the data and wrote the paper. All authors approved the manuscript.

## SUPPLEMENTARY MATERIALS

**Supplementary Figure 1:**
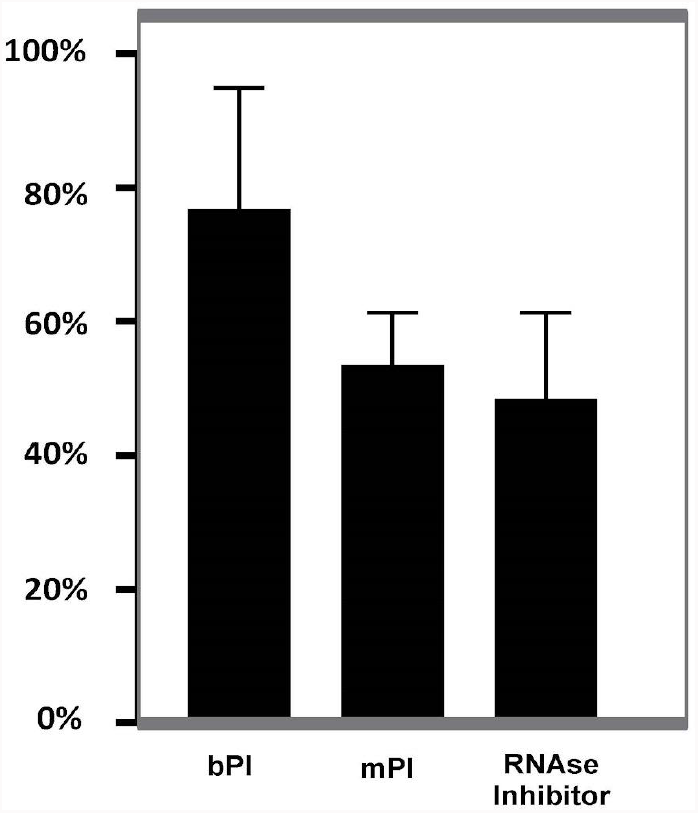
Constitutive sGFP production in absence of clinical samples is diminished by the addition of commercial enzyme inhibitors (mammalian protease inhibitor, bacterial protease inhibitor, and RNase inhibitor). Cell-free reactions were run at 37°C in a sealed 384-well plate for 120 minutes in the presence or absence of RNAse inhibitor, mammalian protease inhibitor, or bacterial protease inhibitor. Data were normalized to a control with no inhibitor and are the mean of three experiments performed on three different days. Error bars correspond to ±SD.

**Supplementary Figure 2:**
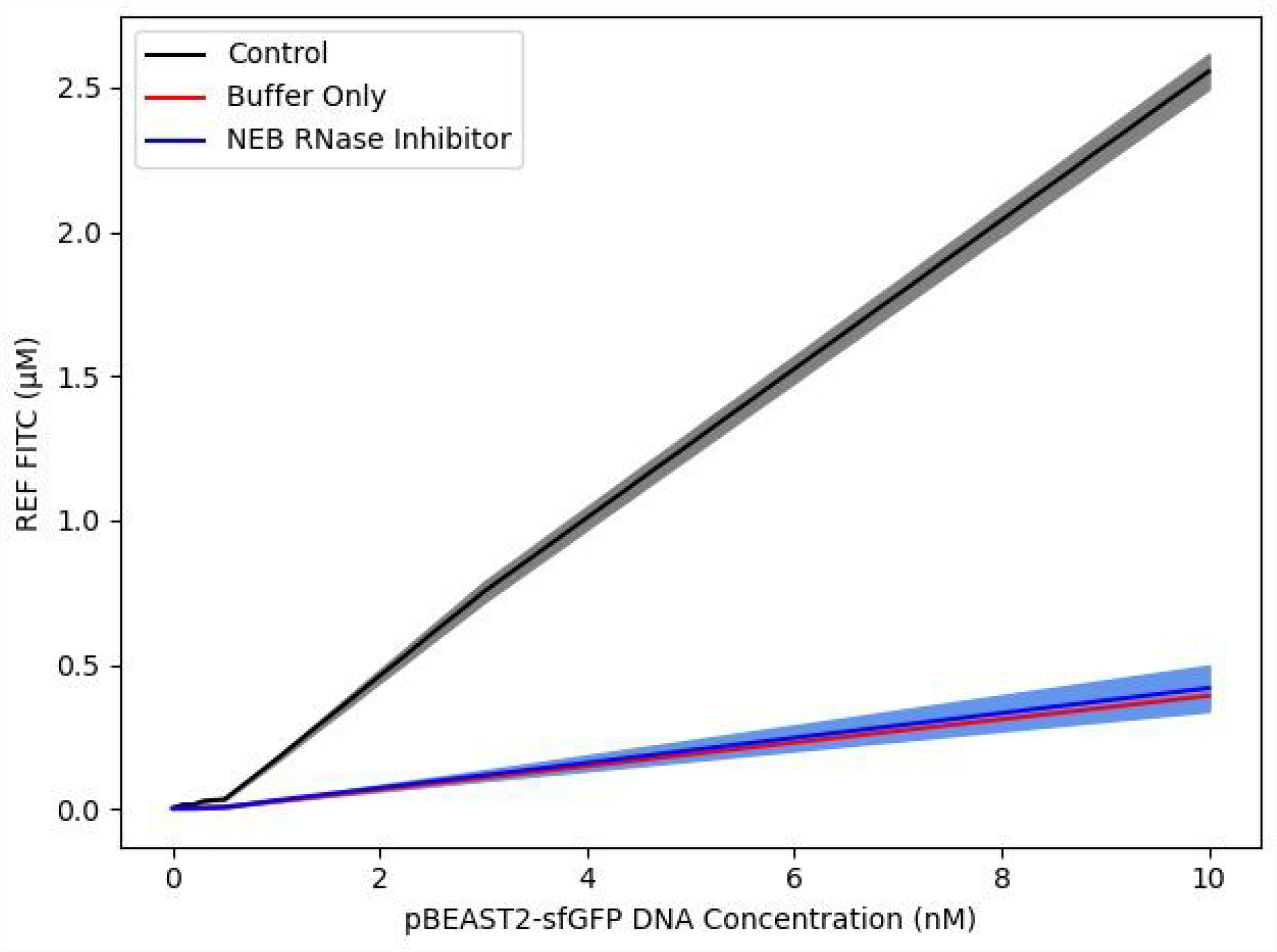
sfGFP expression vs. constitutive DNA concentrations (0-10 nM) for the positive control (standard cell-free reaction mix) and in the presence of RNase inhibitor or equivalent buffer concentration without purified protein (50 mM KCl, 20 mM HEPES, 8 mM DTT, 50% glycerol). Data are the mean of three experiments performed on three different days. Shaded areas correspond to ±SD.

**Supplementary Figure 3:**
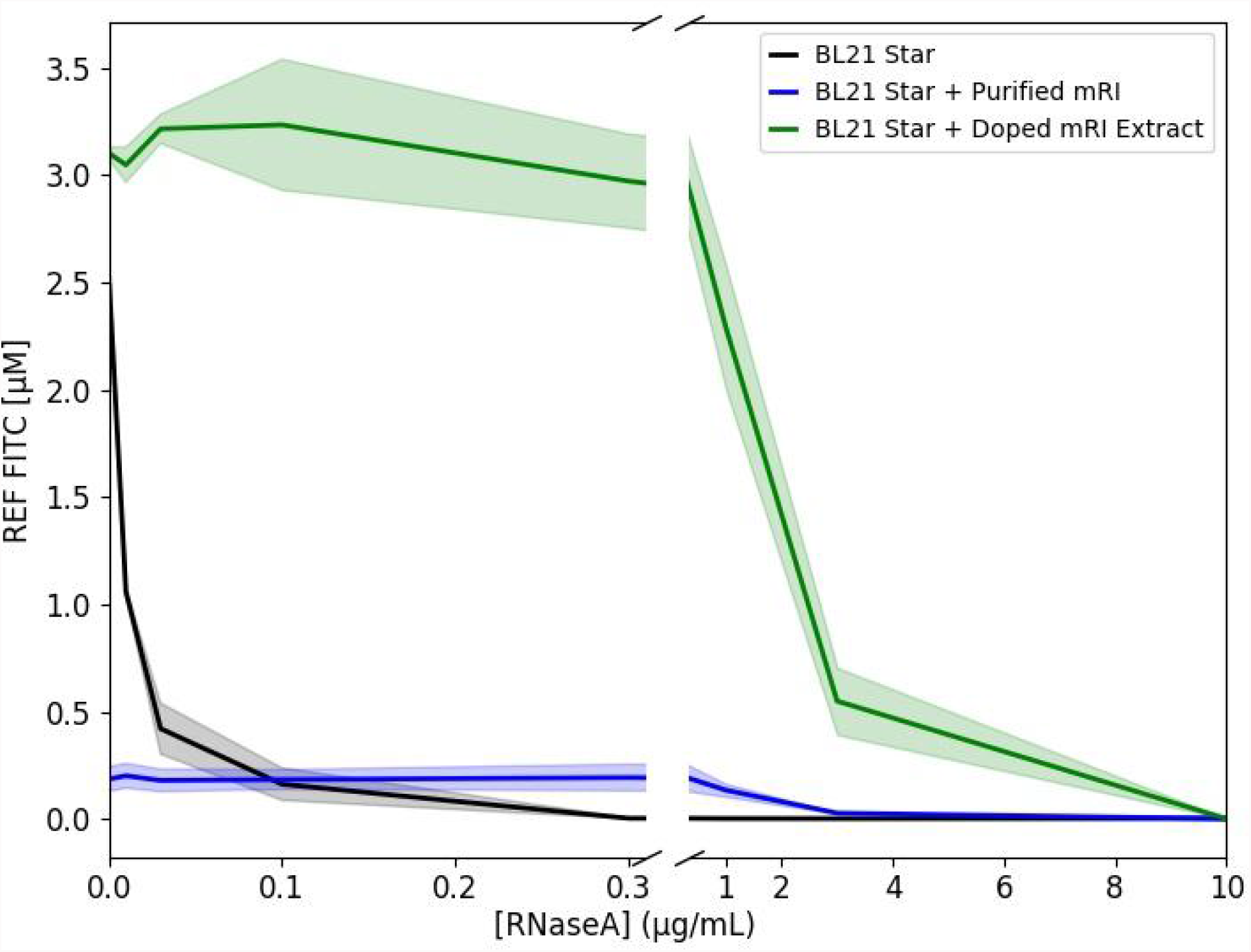
Absolute sfGFP fluorescence (in FITC units) in standard extract (BL21 Star), standard extract with commercial RNase inhibitor (BL21 Star + Purified mRI), and extract induced to produce RNase inhibitor during cell growth (BL21 Star + Doped mRI Extract) when challenged with an RNaseA concentration gradient. Data are the mean of three experiments performed on three different days. Shaded areas correspond to ±SD.

